# Direct HPLC Method for Reduced Pyrroloquinoline Quinone in Functional and Biological Matrices

**DOI:** 10.1101/2025.06.02.657534

**Authors:** Nur Syafiqah Mohamad Ishak, Kazuto Ikemoto

**Affiliations:** Niigata Research Laboratory, Mitsubishi Gas Chemical Company, Inc., Niigata, Japan

**Author notes:** Correspondence should be addressed to: 182, Tayuhama, Kita-ku, Niigata-city, Niigata 950-3112, Japan.

**Keywords:** Pyrroloquinoline quinone (PQQ), Reduced PQQ (PQQ_red_), HPLC, cyclodextrin, ascorbic acid, beverage analysis, redox

## Abstract

Pyrroloquinoline quinone (PQQ) is a redox-active compound with physiological functions, widely used in functional foods and supplements. However, quantifying the reduced form, PQQ_red_, is difficult due to its low solubility and susceptibility to oxidation. This study presents a robust HPLC method for direct quantification of PQQ_red_ in complex matrices. By changing to a strongly acidic eluent, the oxidation of PQQred was suppress, and the optimized method successfully separated PQQred from its oxidized counterpart (PQQ_ox_)and matrix interferences, enabling accurate quantification. A pretreatment using ascorbic acid and γ-CD effectively reduced and solubilized PQQ_red_, and total PQQ can be analyzed as PQQ_red._ The method achieved excellent linearity (R^2^ = 1), low detection limits (LOD: 0.20 mg/L, LOQ: 0.50 mg/L), and high precision (RSD < 3%). Application to commercial beverages showed consistent recovery (99–101%) with minimal interference. Moreover, the method suctandem mass spectrometry.cessfully detected biologically generated PQQ_red_ in yeast cultures, demonstrating its utility in physiological systems. This redox-specific and practical approach enables routine analysis and quality control of PQQ-enriched products, especially where accurate assessment of redox state is essential.

## 1. Introduction

Pyrroloquinoline quinone (PQQ) is a redox-active o-quinone compound that has attracted increasing attention because of its diverse physiological roles including mitochondrial biogenesis, antioxidant defense, cognitive protection, and potential benefits in obesity management [1–5]. As a result, PQQ has gained prominence as a functional food ingredient and is being increasingly incorporated into nutritional supplements and fortified food and beverage products.

PQQ is generally found in an oxidized quinone structure (PQQ_ox_, Fig. 1A). Its reduced counterpart is a quinolinol structure resulting from a two-electron reduction of the quinone moiety (PQQ_red_, Fig. 1B) [6]. The quinone structure demonstrates high reactivity and participates in the acetalization, aldol, and reduction reactions. Furthermore, it irreversibly forms imidazopyrroloquinoline quinones when interacting with amino acids [7,8]. Conversely, the reduced form can coexist with amino acids without undergoing a reaction if it remains unoxidized [9].

**Fig. 1.**
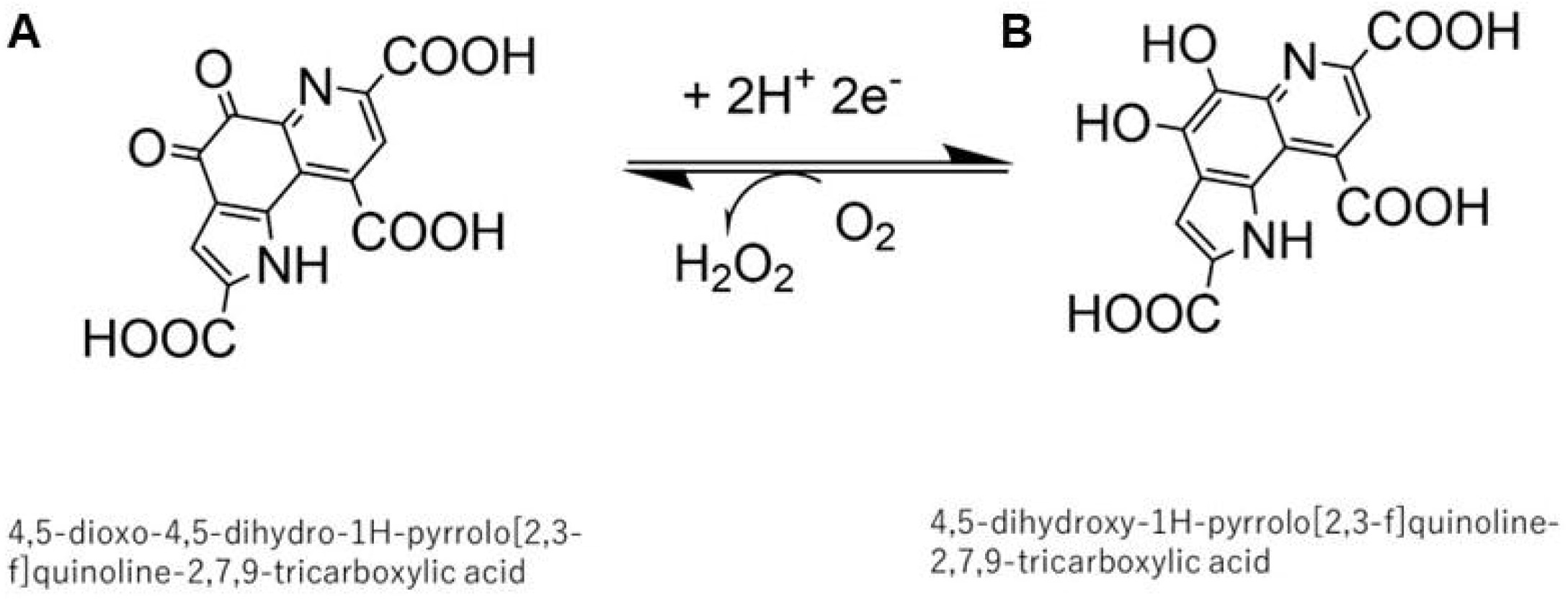
Structural comparison of oxidized and reduced forms of pyrroloquinoline quinone (PQQ). The left structure represents the oxidized form of PQQ (PQQ_ox_), which contains a fully conjugated quinone ring with carbonyl groups at key redox-active positions (A). The right structure illustrates the reduced form (PQQ_red_), in which the quinone ring has been converted into a hydroquinone-like structure through the addition of hydrogen atoms, resulting in hydroxyl groups at the redox-active sites (B).

PQQ_red_ exhibited a strong antioxidant activity [10,11]. However, the quantification of PQQ_red_ is analytically challenging because of its chemical instability and high susceptibility to oxidation. It is typically generated from its oxidized form using ascorbic acid, which is a widely used antioxidant and nutritional additive in food products [12]. Therefore, the analysis of total PQQ was hindered by ascorbic acid. In previous studies, PQQ_red_ has been indirectly measured by converting it to its oxidized form through air oxidation prior to analysis [13] or by applying specialized detection techniques such as colorimetric reactions [14]. However, these approaches are unsuitable for samples rich in reducing agents, such as fortified beverages and functional food products, and often involve complex and time-consuming procedures, which are impractical for routine quality control. Consequently, most existing analytical methods quantify only the oxidized form of PQQ, overlooking the biologically active reduced form and failing to reflect its true concentration or stability in real-world products and biological environments.

To address this limitation, there is a need for a simple, robust, and redox-specific method to quantify the PQQ_red_. Ideally, such a method should be compatible with routine HPLC-UV systems and avoid the use of complex derivatization or electrochemical detection setups. In this study, we present a validated HPLC method that enables direct detection and quantification of PQQ_red_. This approach addresses the key challenges in PQQ_red_ analysis and offers a practical solution for both research and quality control applications.

## 2. Results and Discussion

### 2.1. Separation of PQQ Redox Forms by HPLC

In conventional analyses using weakly acidic eluents [13], oxidized and reduced forms of PQQ are not well separated, and PQQ_red_ often undergoes oxidation during analysis. Additionally, ascorbic acid—commonly used as a reducing agent—elutes near the PQQ peaks, leading to interference and reduced specificity (Fig. 2A).

**Fig. 2.**
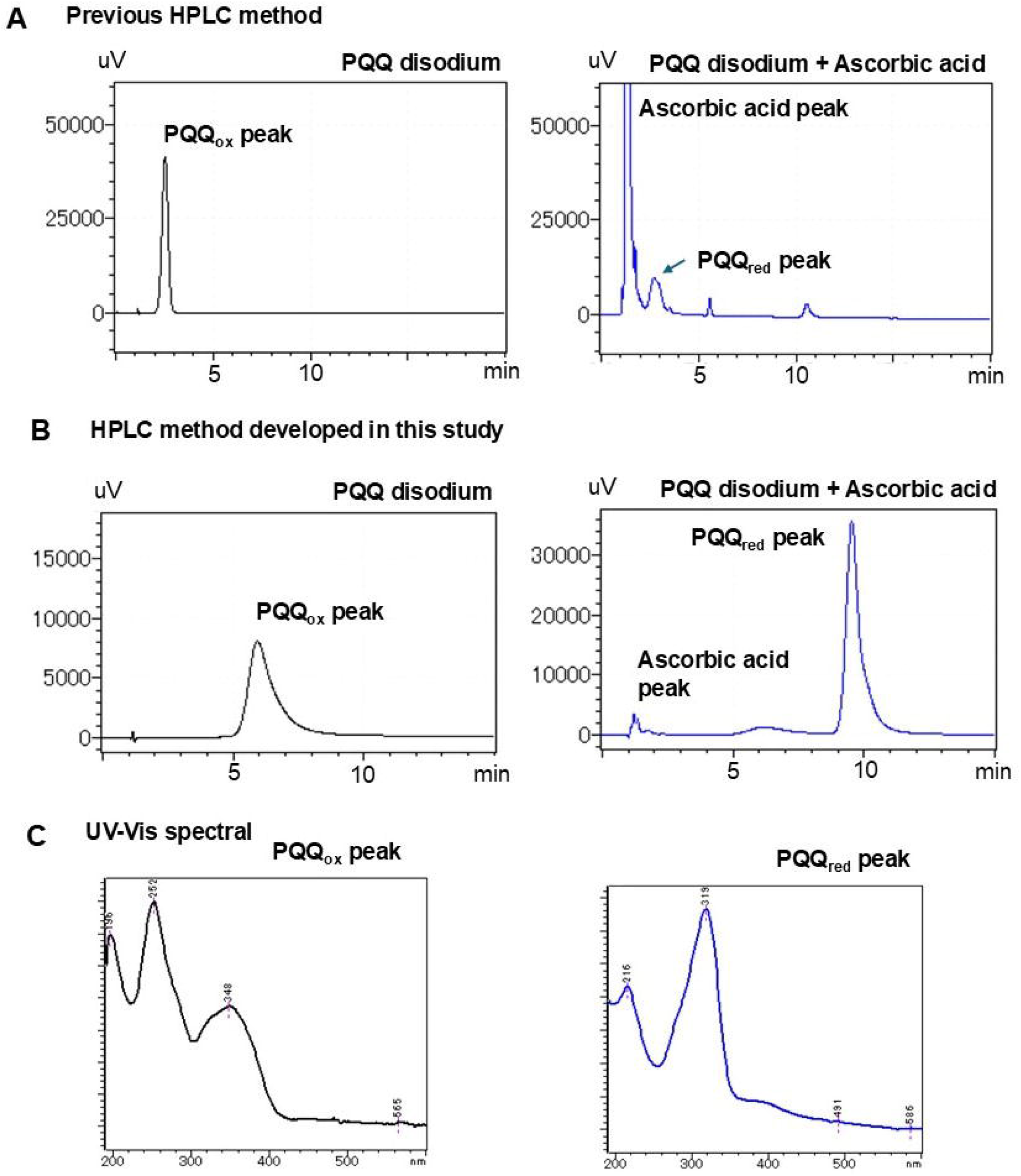
Chromatographic separation and UV-visible spectra of PQQ_ox_ and PQQ_red_. Representative HPLC chromatograms from the previous method (A) and the method developed in this study (B) are shown. The left panel of each chromatogram shows the result for the PQQ_ox_ standard (∼2.5 min), while the right panel shows the chromatogram following the reduction of PQQ (∼2.8 min) by ascorbic acid. In (B), the distinct separation of PQQ_red_ (∼10 min) from PQQ_ox_ (∼6 min) was obtained. UV-visible absorption spectra (C) of the PQQ_ox_ and PQQ_red_ peaks from chromatogram (B) confirm their redox states: PQQ_ox_ exhibits a maximum absorbance at 252 nm, whereas PQQ_red_ shows a distinct peak at 318 nm.

To address these limitations, our strategy involves in using a strongly acidic mobile phase, which stabilizes PQQ_red_ by suppressing its oxidation and proton dissociation, thus enhancing the physical property distinction between PQQ_ox_ and PQQ_red_. This approach allowed successful separation on an ODS column. Under these optimized conditions, PQQ_red_ eluted as a sharp peak at approximately 10 minutes, while PQQ_ox_ appeared at around 6 minutes, enabling clear separation on an ODS column (Fig. 2B). The identity of each peak was confirmed by their distinct UV-Vis spectra: PQQ_ox_ at ∼250 nm and PQQ_red_ at ∼320 nm (Fig. 2C). Ascorbic acid eluted much earlier than both PQQ species, with no co-elution observed from matrix components, confirming the specificity of the method.

### 2.2. Improving the Solubility and Stability of PQQ_red_

For total PQQ quantification, we aimed to convert all PQQ into its reduced form (PQQ_red_) prior to HPLC analysis. This strategy allows for a single calibration curve to simplify the quantification. It also avoids unwanted side reactions, as the quinone structure in PQQ_ox_ is reactive toward nucleophilic compounds such as amines. However, PQQ_red_ is chemically unstable in weakly acidic conditions, making strongly acidic environments essential for preserving its reduced state during sample preparation and chromatographic analysis. Therefore, we developed pretreatment process designed to convert and stabilize all PQQ as its reduced form. PQQ_red_ exhibits low aqueous solubility, particularly at low pH. Our preliminary solubility tests indicated that solubility drops markedly below pH 3.8, with measured values of 12, 14, and 50 mg/L at pH 2.8, 3.1, and 3.4, respectively (Fig. 3A). This was also evident during reduction with ascorbic acid, where visible precipitation of PQQ_red_ was frequently observed, indicating poor solubility in acidic environments.

**Fig. 3.**
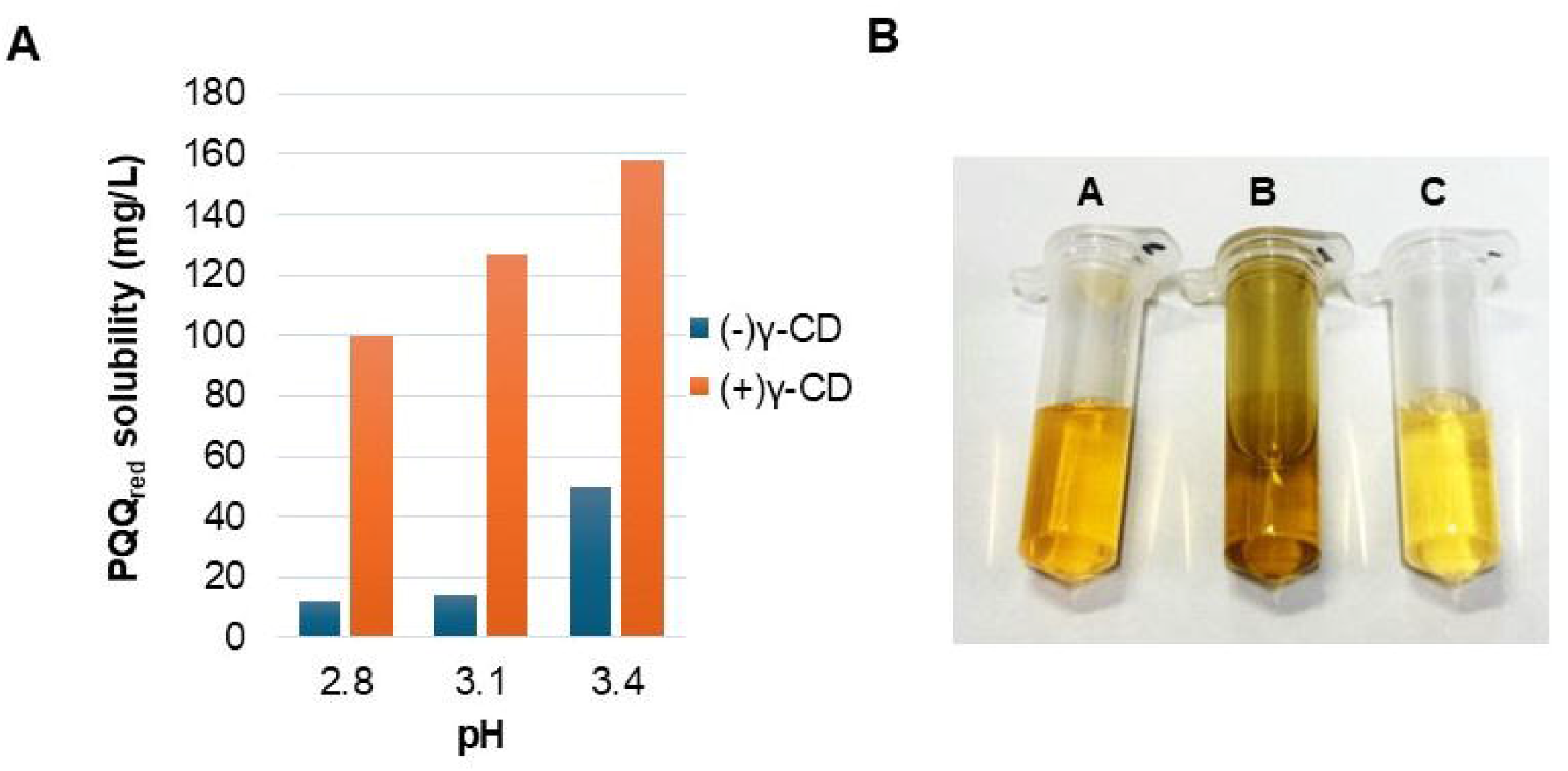
Effect of pH and γ-CD on the solubility of PQQ_red_. Solubility of PQQ_red_ was measured at three pH values (2.8, 3.1, and 3.4) in the absence [(−) γ-CD] and presence [(+) γ-CD] of 10% (w/w) γ-CD (A). The addition of γ-CD significantly enhanced PQQ_red_ solubility in acidic pH levels. Representative images of PQQ solutions after reduction under different conditions (B): Tube A –PQQ treated with 5% ascorbic acid and 10% γ-CD (clear solution), Tube B –PQQ treated with 5% ascorbic acid only (showing precipitation), Tube C –PQQ in phosphate buffer (control).

To overcome this, we evaluated γ-cyclodextrin (γ-CD), a compound widely used in foods to enhance solubility through inclusion complex formation [15,16]. The addition of 10% (w/w) γ-CD to the pretreatment solution significantly improved solubility, increasing PQQ_red_ concentrations to 100, 127, and 158 mg/L at pH 2.8, 3.1, and 3.4, respectively, and effectively preventing precipitation (Fig. 3B). Based on these findings, we established a pretreatment protocol using 5% (w/w) ascorbic acid and 10% (w/w) γ-CD. This combination achieved nearly complete recovery (100%) and excellent stability after 24 hours (101%), as shown in Table 1. In comparison, ascorbic acid alone resulted in only 11% initial recovery and 3% after 24 hours, while the untreated control showed no reduction and no detectable PQQ_red_.

**Table 1.**
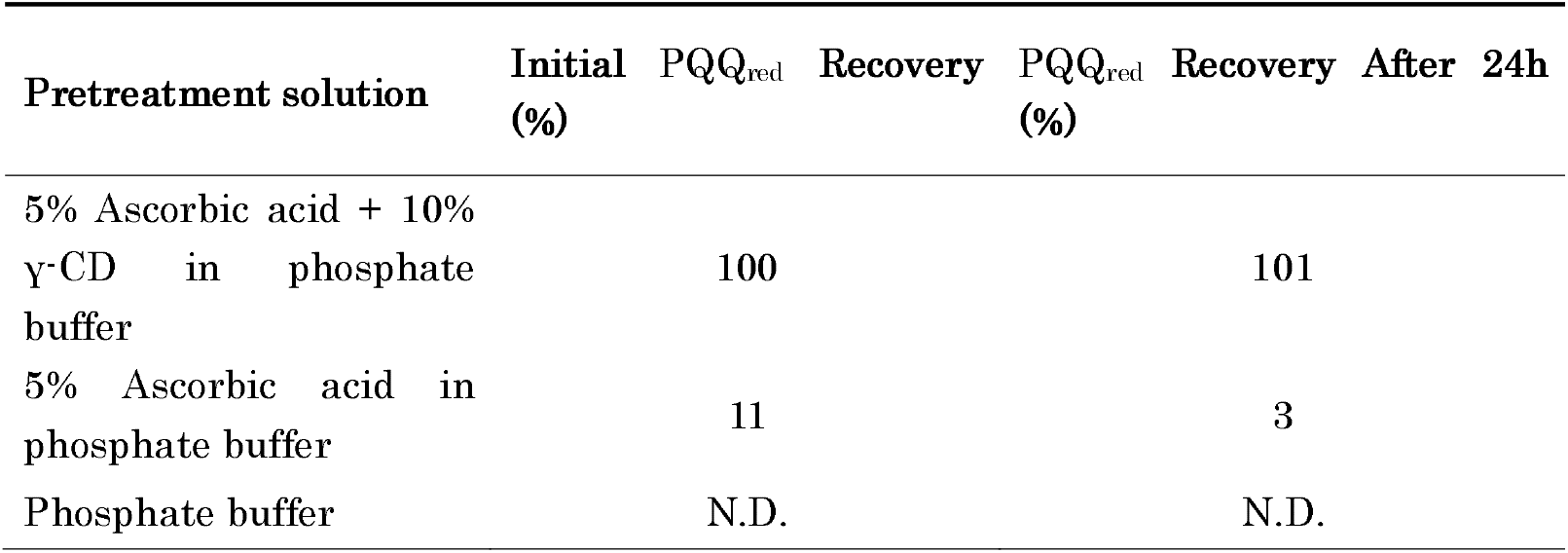
Stabilization of PQQ_red_ under different pretreatment conditions. The recovery of was evaluated immediately after pretreatment (Initial recovery) and after 24 h at room temperature. N.D.: Not detected.

### 2.3. Linearity and Quantification Limits

To evaluate the quantitative performance of the method, a calibration curve was constructed using PQQ disodium standard solutions ranging from 2 to 80 mg/L. Each solution was treated in a 1:1 ratio with a pretreatment mixture containing 5% (w/w) ascorbic acid and 10% (w/w) γ-CD to ensure high stability and solubility of PQQ_red_. The resulting calibration curve exhibited excellent linearity with a regression equation of y = 13,514x – 27,796 and a correlation coefficient of R^2^ = 1 (Fig. 4A), confirming the suitability of the method for precise quantification.

**Fig. 4.**
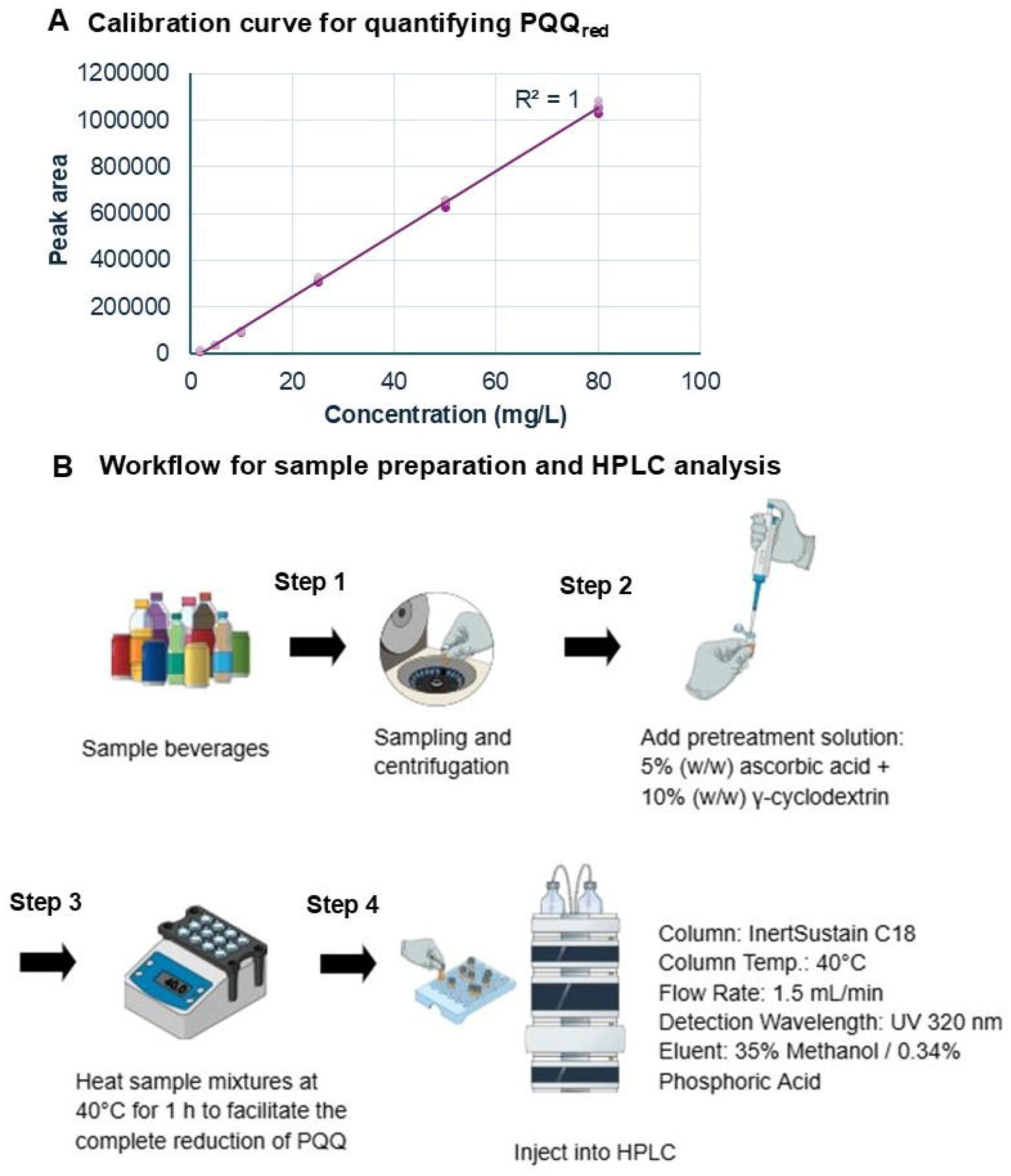
Calibration curve (A) and sample preparation workflow for the quantification of PQQ_red_ using HPLC (B). The calibration curve was constructed using PQQ disodium standards treated with pretreatment solution, showing excellent linearity for the range of 2–80 mg/L. Sample preparation steps: Step 1 – Commercial beverage samples were centrifuged to remove particulates. When the PQQ disodium concentration exceeded the calibration range, the samples were diluted with phosphate buffer prior to pretreatment. Step 2 – A pretreatment solution containing 5% (w/w) ascorbic acid, and 10% (w/w) γ-CD was added. Step 3 – The mixture was incubated at 40°C for 1 hour to facilitate the complete reduction of PQQ_ox_ to PQQ_red_. Step 4 – The samples were injected into an HPLC system for analysis under optimized conditions.

The relative standard deviation (RSD) values across the tested concentrations were consistently below 3%, ranging from 2.63% to 2.96%. Only the lowest level (2 mg/L) showed a slightly higher RSD of 4.81%, which was considered acceptable because of the greater variability near the detection limit. The limit of detection (LOD) and limit of quantification (LOQ), calculated based on the standard deviation at 2 mg/L and the slope of the calibration curve, were 0.20 mg/L and 0.50 mg/L, respectively. These results confirm that the method provides high sensitivity, accuracy, and reproducibility, making it suitable for the quantitative analysis of PQQ_red_ in complex sample matrices such as functional beverages and dietary supplements.

### 2.4 Quantification of PQQ_red_ in Commercial Beverages

To assess the robustness and real-world applicability of this method, the protocol was applied to beverages spiked with known amounts of PQQ. The sample preparation and HPLC analysis workflow is exhibited in Fig. 4B. Samples A and B were analyzed in triplicate, while samples C–F were measured once. The measured PQQ_red_ concentrations in samples A and B were consistent with the spiked values, yielding recovery rates of 100% and 99%, respectively (Table 2). The method demonstrated excellent repeatability, with RSD values of 0.3% for sample A and 0.9% for sample B, indicating robust performance, even in complex beverage compositions. In samples C–F, recoveries ranged from 100% to 101%, further supporting the reliability and compatibility of the method with a variety of beverage ingredients, including those containing high levels of ascorbic acid, citric acid, or sweeteners (Table 2). No matrix interference or peak distortion was observed, confirming the specificity of this method. These results validated the suitability of this method for routine quality control and label verification of PQQ-containing functional beverages.

**Table 2.**
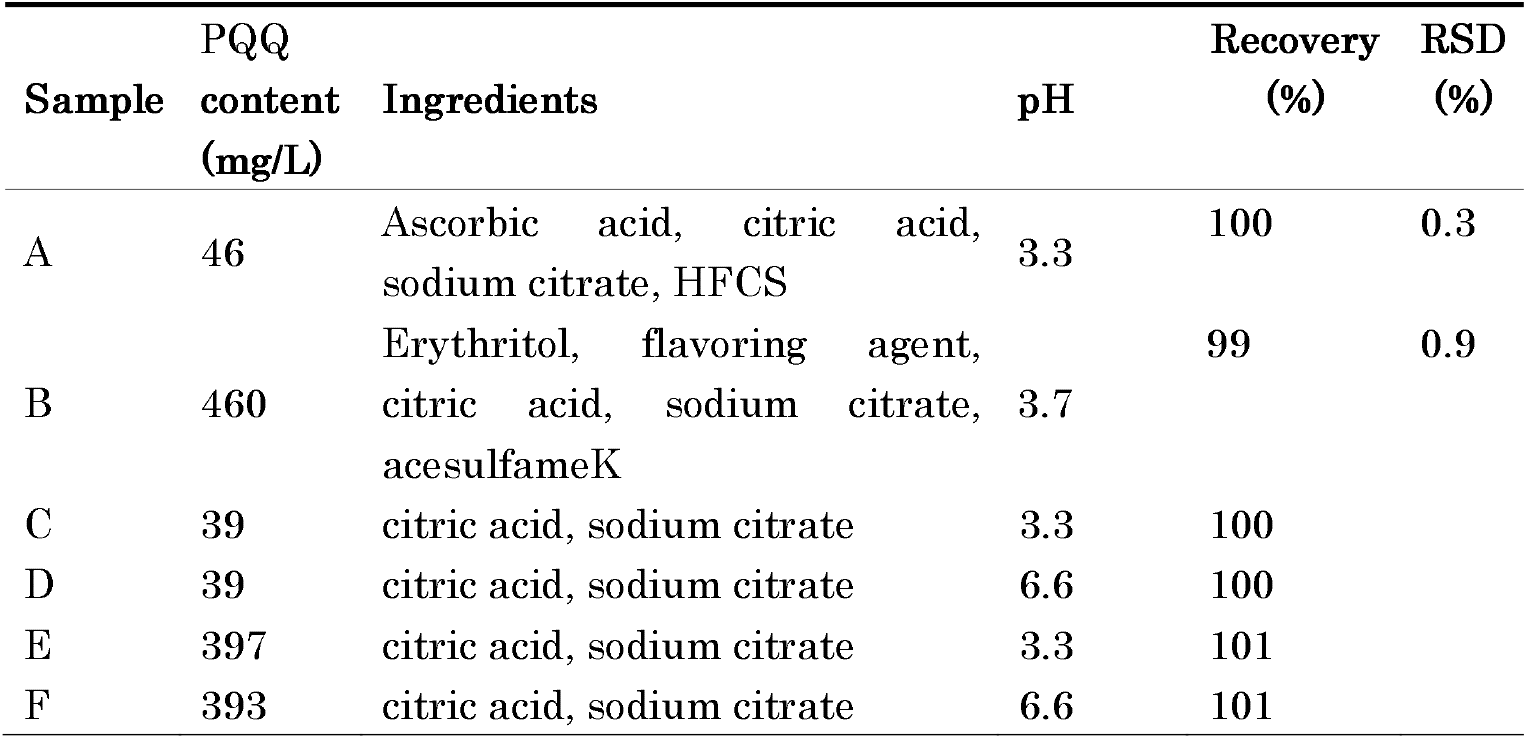
Measured concentrations of PQQ_red_ in spiked commercial beverage samples using the developed HPLC method. Samples A and B were analyzed in triplicate and include statistical parameters: average concentration, standard deviation (SD), relative standard deviation (RSD%), and recovery rate. Samples C–F were analyzed as single measurements.

| Sample | PQQ content (mg/L) | Ingredients | pH | Recovery (%) | RSD (%) |
| --- | --- | --- | --- | --- | --- |
| A | 46 | Ascorbic acid, citric acid, sodium citrate, HFCS | 3.3 | 100 | 0.3 |
| B | 460 | Erythritol, flavoring agent, citric acid, sodium citrate, acesulfameK | 3.7 | 99 | 0.9 |
| C | 39 | citric acid, sodium citrate | 3.3 | 100 |  |
| D | 39 | citric acid, sodium citrate | 6.6 | 100 |  |
| E | 397 | citric acid, sodium citrate | 3.3 | 101 |  |
| F | 393 | citric acid, sodium citrate | 6.6 | 101 |  |

### 2.5. Detection of biologically reduced PQQ in yeast culture

To investigate whether biologically generated PQQ_red_ can be detected using this develop HPLC method, a simple experiment was conducted using a yeast culture. PQQ was added to *S. cerevisiae* in glucose-containing solution and incubated at 28°C for 20 h. As shown in Fig. 5, a noticeable color change occurred in the yeast-containing solution. HPLC analysis revealed a distinct peak corresponding to PQQ_red_, alongside the PQQ_ox_ peak. In contrast, the control sample—containing PQQ and glucose but without yeast—showed only a minor PQQ_red_ signal. These findings demonstrate that the method is capable detecting PQQ_red_ formed under biological conditions. The quantified amount of PQQ_ox_ and PQQ_red_ in both samples corresponding to HPLC peak areas is summarized in Table 3.

**Table 3.** The amount of PQQ_ox_ and PQQ_red_ calculated from HPLC peak areas in samples incubated with and without yeast.

| Sample | PQQ<sub>ox</sub> amount (mg/L) | PQQ<sub>red</sub> amount (mg/L) |
| --- | --- | --- |
| Yeast + Glucose + BioPQQ | 102.1 | 33.9 |
| Glucose + BioPQQ | 162.2 | 4.4 |

**Fig. 5.**
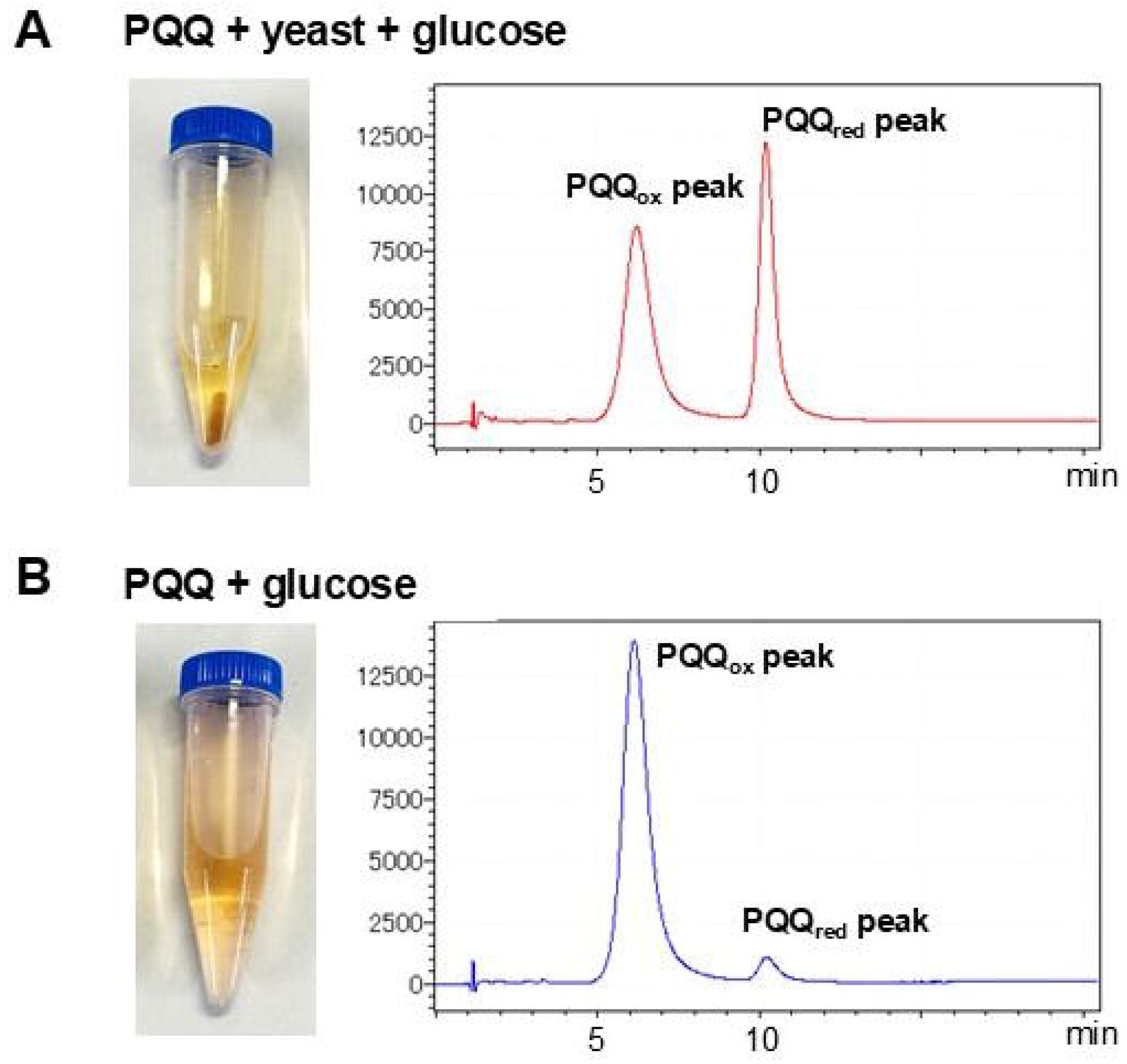
Detection of PQQ_red_ in a biological environment using the developed HPLC method. Representative chromatograms and corresponding test tube images of samples containing 2 g/L PQQ disodium with glucose and glucose (A) and without yeast (B), after incubation at 28°C for 20 hours. In the sample with yeast, a distinct PQQ_red_ peak was observed alongside PQQ_ox_, indicating that PQQ_ox_ was reduced under biological conditions. In contrast, the control sample (without yeast) showed a much smaller PQQ_red_, reflecting limited reduction.

## 3. Discussion

This study successfully established a direct and robust HPLC method for the quantification of PQQ_red_ in complex matrices using a strongly acidic eluent. Unlike conventional approaches, which often use weakly acidic mobile phases and cannot achieve chromatographic separation between oxidized and reduced forms, the present method leverages strong acidity to suppress oxidation of PQQ_red_ and reduce proton dissociation [17,18]. This enhanced the physicochemical differences between quinone and hydroquinone structures, enabling their effective separation on an ODS column. In traditional method [9,13], the elution of PQQ closely overlapped with that of ascorbic acid, complicating analysis in real-world matrices like beverages. In contrast, our optimized method achieved clear separation, with PQQ_ox_ eluting at ∼6 min and PQQ_red_ at ∼10 min, and early elution of ascorbic acid, minimizing interference.

To analyze total PQQ as its reduced form, we designed a pretreatment protocol combining ascorbic acid as a reducing agent with γ-CD to improve solubility. This was crucial since PQQ_red_ is poorly soluble at low pH and prone to precipitation. Our solubility tests demonstrated that γ-CD significantly increased solubility at acidic pH, consistent with pharmaceutical and food applications of γ-CD to enhance aqueous solubility through inclusion complexation [16,19]. The combined pretreatment (5% ascorbic acid and 10% γ-CD) yielded nearly complete recovery and excellent stability, outperforming ascorbic acid alone.

Application to commercial beverages confirmed the method’s practicality. Spiked samples showed recoveries of 99–101% with RSD <1%, and no observable matrix interference, even in products containing high levels of acids or sweeteners. This confirms the method’s suitability for routine quality control in functional beverages and supplements. The validated method meets AOAC and USP analytical performance standards [20–22], including excellent linearity (R^2^ = 1), low LOD/LOQ (0.20 / 0.50 mg/L), and precision (RSD < 3% at all but the lowest concentration). It thus fulfills key criteria for food and supplement analysis and is suitable for regulatory compliance and industrial applications.

While advanced techniques such as LC-MS/MS [23,24] and enzymatic assays [14] have been developed for PQQ quantification in food matrices, they often require specialized instrumentation, prolonged sample preparation, and may not maintain the redox state of PQQ during analysis. These methods, although highly sensitive, are less practical for routine quality control or manufacturing settings. In addition, PQQ has been analyzed by derivatizing with amino acids (e.g., glycine) to form imidazolopyrroloquinoline (IPQ) derivatives[9]. While this derivatization method provides high sensitivity and selectivity for PQQ_ox_, it is unsuitable for quantifying PQQ_red_, as the condensation reaction depends on electrophilic carbonyl groups present only in the quinone structure. Moreover, the method is incompatible with matrices containing excess reducing agents, which inhibit the derivatization reaction by maintaining PQQ in its reduced state. In contrast, the method developed in this study employs a simple reverse-phase HPLC-UV system that is accessible and cost-effective for many laboratories. Importantly, it is specifically designed to preserve and quantify the biologically active PQQ_red_, which is not targeted by most existing protocols.

Notably, this method enabled simultaneous detection of PQQ_ox_ and PQQ_red_ in a biological environment. Yeast cultures incubated with PQQ and glucose produced measurable levels of PQQ_red_, suggesting *in vivo* reduction. The ability to directly quantify redox states represents a significant advancement in redox biology. This technique provides new insights into the redox dynamics of PQQ and may aid in evaluating bioavailability, redox cycling, and antioxidant function *in vivo*. Such capability is essential for evaluating the effectiveness of PQQ supplementation and functional foods, as well as for advancing redox biology research through potential biomarker development akin to GSH/GSSG or NAD^+^ /NADH ratios [25]. The current method addresses this gap by allowing accurate and redox-sensitive quantification in complex biological matrices, advancing the field of redox biology and antioxidant research.

Beyond food analysis, the method opens new avenues in biological and engineering research. For example, our findings suggest that yeast can reduce extracellular PQQ, with implications for cellular redox control and even bioenergy applications. Suppose a turbidity of 1 is 10^8^ cell, one yeast cell is reducing 6 × 10^11^ molecules of PQQ. The reduction of extracellular PQQ suggests that its distribution within the body may be heavily skewed toward the reduced form, indicating a strong potential for reduction in the bloodstream. The observed minor reduction of PQQ in the glucose-only control suggests intrinsic catalytic behavior of PQQ in redox reactions, possibly enabling new catalytic or bio-electrochemical applications. From an engineering perspective, this discovery opens possibilities for utilizing cells as mediators to extract energy. Specifically, it may enhance the efficiency of charge extraction in microbial fuel cells. Alternatively, biological systems could be engineered to produce reduced PQQ for energy storage applications.

## 4. Materials and Methods

### 4.1. Chemicals and reagents

Pyrroloquinoline quinone (PQQ) used in this study was the disodium salt trihydrate form (PQQ·Na_2_·3H_2_O), commercially known as BioPQQ®, supplied by Mitsubishi Gas Chemical Co., Ltd. Ascorbic acid, γ-cyclodextrin (γ-CD), phosphoric acid (85%), hydrochloric acid (36%), methanol (HPLC grade), and acetonitrile (HPLC grade) were purchased from Fujifilm Wako Pure Chemical Industries, Japan. Ultrapure water was prepared using a Milli-Q purification system (Millipore, USA) and was used for all solution preparations and HPLC procedures.

Pure PQQ_red_ standard was prepared using a modified reduction protocol. A solution of 3.0 g PQQ disodium salt in 1.2 L of water was added to a mixture containing 30 g L-ascorbic acid, 120 g water, and 2.5 g of 2 N hydrochloric acid, maintained at 12°C. The mixture was stirred at 20°C for 20 hours, followed by the addition of another 2.5 g of 2 N HCl. The resulting crystals were filtered, washed with 5 mL of 2 N HCl and 8 mL of 50% ethanol, and dried under vacuum at room temperature for 20 hours. This yielded 3.35 g of yellow brown PQQ_red_ crystal.

### 4.2. Chromatographic conditions

High-performance liquid chromatography (HPLC) analysis was performed using a system equipped with a UV-Vis detector set at 320 nm. A C18 reverse-phase column (InertSustain C18, 4.6 × 15 cm, 5 μm particle size) was employed as the stationary phase. The mobile phase consisted of 0.34% (w/w) phosphoric acid and 35% (v/v) methanol in water (pH2.5). The flow rate was set at 1.5 mL/min, and the column temperature was maintained at 40°C. The injection volume for all analyses was 10 μL. For comparison, a prior HPLC method employed a system with a UV detector set at 259 nm. A C18 reverse-phase column (YMC-Pack ODS-A, 4.6 × 15 cm, 5 μm particle size) was utilized as the stationary phase. The mobile phase was composed of a 30:70 (v/v) mixture of 100 mM acetic acid and 100 mM ammonium acetate, adjusted to pH 5.1. The flow rate was similarly set at 1.5 mL/min, with the column temperature maintained at 40°C. The injection volume in this method was 10 μL.

### 4.3. Preparation of PQQ standard solutions

A stock solution of PQQ disodium (1 mg/mL) was prepared in distilled water. For calibration, working standard solutions were freshly prepared by serial dilution with water or phosphate buffer immediately prior to the analysis.

### 4.4. Evaluation of pre-treatment solution

To assess the effectiveness of the pretreatment solution in stabilizing the PQQ_red_, several conditions were compared using aqueous PQQ_ox_ standards. The tested pre-treatment formulations included 5% (w/w) ascorbic + 10% (w/w) γ-cyclodextrin acid; 5% (w/w) ascorbic acid only; and Control buffer. All solutions were prepared in a 3.4% phosphoric acid solution. Each formulation was mixed 1:1 with a 0.1 mg/L PQQ_ox_ solution and incubated at 40°C for 1 h. After the treatment, the solutions were cooled to room temperature and centrifuged for 1 min. The samples were filtered, transferred into HPLC vials, and immediately analyzed using HPLC. The peak area of the PQQ_red_ was used as a measure of the stabilization efficiency. Stability was also evaluated after 24 h at room temperature to assess the retention of the PQQ_red_ post-treatment. The recovery percentage was calculated by normalizing the peak area to the initial highest value observed under the tested conditions, which was defined as 100% recovery.

### 4.6. Quantification and calibration

The PQQ_red_ was quantified using a freshly prepared external standard solution, prepared by reducing PQQ_ox_ with excess ascorbic acid under the same conditions used for sample pretreatment to ensure matrix consistency. Calibration curves were constructed by plotting the peak area against known concentrations of the PQQ_red_ standards (2–80 mg/L), each analyzed in triplicate. The LOD and LOQ for PQQ_red_ were estimated based on the standard deviation of the response and the slope of the calibration curve. Using the standard deviation of the peak areas at the lowest tested concentration (2 mg/L) and the slope calculated from the linear portion of the curve, LOD and LOQ were calculated as follows: LOD= (3.3 × σ)/slope, LOQ = (10 × σ)/slope.

### 4.5. Samples preparation

The sample beverages were selected as model matrices for analysis. The beverage samples were spiked with known amounts of PQQ (Table 4). Samples with PQQ concentrations exceeding 80 mg/L were diluted in the 5–80 mg/L range to prevent the precipitation of PQQ_red_, which tends to occur at higher concentrations owing to its limited solubility. Prior to analysis, the beverages were degassed by mild sonication and centrifuged to remove the particulate matter. A pre-treatment solution containing 5% (w/w) ascorbic acid and 10% (w/w) γ-cyclodextrin in phosphate buffer was added to the sample solution at a 1:1 ratio. The mixture was incubated at 40°C for at least 1 h to ensure complete reduction of PQQ. The prepared samples were immediately injected into the HPLC system.

### 4.6. Detection of Biologically PQQ_red_ in Yeast Culture

A model experiment using *Saccharomyces cerevisiae* NBRC 100929 was conducted to evaluate whether the developed method could detect biologically PQQ_red_. First, a yeast suspension was prepared by inoculating 5 mL of autoclaved YEPD medium (containing 10 g/L yeast extract, 20 g/L Bacto peptone, and 20 g/L glucose) with a glycerol stock of *S. cerevisiae*. The culture was incubated overnight at 28°C. Following cultivation, the yeast cells were harvested by centrifugation, and the supernatant was discarded. The cell pellet was resuspended in 2% (w/v) glucose solution to adjust the optical density to OD600 = 12.

For the PQQ reduction reaction, 0.5 mL of PQQ solution (2 g/L in phosphate-buffered saline, PBS; pH 7.2), 5.0 mL of 2% glucose solution, and 0.5 mL of the yeast suspension were combined in a test tube. The mixture was incubated at 28°C for 20 hours with shaking. After incubation, the reaction mixture was centrifuged to remove yeast cells, and the supernatant was collected for HPLC analysis.

A control sample was prepared by mixing 0.5 mL of PQQ disodium solution (2 g/L in PBS, pH 7.2) with 5.5 mL of 2% glucose solution, excluding yeast. This mixture was incubated under the same conditions (28°C, 20 hours, shaking) and processed identically. The presence of PQQ_red_ was then assessed using the developed HPLC method.

## 5. Conclusion

This study offers a sensitive, reliable, and redox-specific tool for the direct quantification of PQQ_red_ in complex matrices. The simplifies method exhibited excellent linearity, precision, and sensitivity, and was successfully applied to spiked commercial products. Furthermore, the HPLC condition developed in this study demonstrated the capability to detect both oxidized and reduced forms of PQQ in a biological environment, highlighting its versatility and relevance for both nutritional and biological contexts. Overall, the approach provides a redox-specific, user-friendly analytical tool suitable for routine analysis and quality control in both research and industrial settings, particularly where the accurate evaluation of the PQQ redox state is essential.

## Abbreviations

PQQ: Pyrroloquinoline quinone
HPLC: High Performance Liquid Chromatography
UV: Ultraviolet radiation
γ-CD: gamma-cyclodextrin
LOD: Limit of detection
LOQ: Limit of quantification
RSD: Relative standard deviation.

## Author Contributions

NSMI conducted the experiments, analyzed the data, and wrote the manuscript. KI supervised the research design and data interpretation. Both authors contributed to the discussion, reviewed the manuscript, and approved the final version for submission.

